# Patching the Leak or Rebuilding the Boat? Evaluating Targeted Probiotic Cyanobacteria and Microbiome Transplants to Counteract Antibiotic‑Driven Rhizosphere Dysbiosis in Tomato under *Xanthomonas perforans* Pressure

**DOI:** 10.64898/2026.05.20.726701

**Authors:** Toi Ketehouli, Erica M. Goss, Fabiano José Perina, Samuel J. Martins

## Abstract

Antibiotic use in agricultural systems can unintentionally disrupt beneficial rhizosphere microorganisms, yet the consequences of this dysbiosis for plant fitness remain insufficiently understood. Building on previous findings that application of streptomycin to the roots decreases cyanobacteria and increases tomato plant susceptibility to foliar *Xanthomonas* infection, this study aimed to determine whether this relationship reflects causation or correlation. We evaluated whether targeted inoculation with the filamentous nitrogen-fixing cyanobacterium *Cylindrospermum* sp. (CI) or a complex rhizosphere microbiome transplant (RMT) could mitigate antibiotic-induced dysbiosis. As expected, streptomycin treatment significantly increased bacterial spot disease severity and reduced microbial richness in the rhizosphere, marked by a pronounced decline in cyanobacterial and *Cylindrospermum* operational taxonomic units. Co-occurrence network analysis revealed that this dysbiotic state was defined by reduced community connectivity and increased negative associations, indicating a breakdown in cooperative microbial relationships. Notably, both CI and RMT reduced plant disease severity, though they caused distinct rhizosphere community reassembly outcomes. While RMT relied on microbial functional redundancy, the targeted CI approach achieved more robust colonization and effectively “patched” the functional gap left by dysbiosis. Microbiome restoration directly influenced host physiology, significantly reducing the overactivation of ethylene-mediated defense genes, such as *ERF1*, and partially reinstating auxin-responsive signaling pathways (*IAA21*) that were disrupted under dysbiosis. These findings suggest that targeted microbial inoculation could reverse dysbiosis and enhance plant resilience under pathogen pressure as effectively as complex microbial transplants. This work highlights a shift in microbiome management: from the complex ‘rebuilding’ of communities to the strategic ‘repair’ of specific functional gaps.

## Introduction

Tomato (*Solanum lycopersicum* L.) is one of the most economically important vegetable crops worldwide; however, its production is severely constrained by bacterial leaf spot disease, caused by *Xanthomonas perforans* (syn. *X. euvesicatoria* pv. *perforans*). In Florida, bacterial spot disease can reduce marketable yield by up to 66%^1^. After decades of reliance on copper-based bactericides and antibiotics, including the widespread historical use of streptomycin, disease control remains largely inadequate because of pathogen resistance^1,2^. Growing concerns about antimicrobial resistance, phytotoxicity, and microbiome disruption have underscored the need for sustainable disease management strategies that enhance plant immunity rather than suppress pathogens through chemical toxicity^3^. Recent work highlights the importance of the rhizosphere microbiome in supporting plant growth, nutrient cycling, and immune function^4^. Certain plant-associated microbes are essential for plant physiological functions and serve as the initial defense against invading pathogens^5-8^, representing an “extended host immune system.”^9^. Plant defense facilitated by the rhizosphere microbial community can occur directly through the synthesis of antimicrobial compounds and competition for resources, or indirectly by triggering the plant’s defense signaling pathways, such as the induction of systemic resistance (ISR)^10-12^. A disruption of microbial communities can lead to the breakdown of host-microbiota homeostasis, resulting in reduced host fitness, a condition known as dysbiosis.

Borrowing from the success of fecal microbiome transplants in human medicine, rhizosphere microbiome transplant (RMT) has emerged as a strategy to restore plant health. By transferring complex microbial communities from disease-suppressive “donor” soils to “recipient” dysbiotic plants RMT aims to re-establish homeostatic defense mechanisms and suppress pathogen proliferation^13-16^. Our previous research demonstrated that streptomycin-induced dysbiosis depleted beneficial filamentous heterocyst-forming genera, such as *Cylindrospermum* spp., from the cyanobacteria phylum^17^. Cyanobacteria are emerging as key contributors to soil health and plant resilience because of their ability to fix atmospheric nitrogen, produce phytoactive compounds, modulate soil structure, and support microbial community stability^18-20^. *Cylindrospermum* spp. are particularly valuable because they conduct oxygen-sensitive nitrogen fixation, exude bioactive metabolites, and establish beneficial interactions with plant roots^21,22^. These traits position *Cylindrospermum* spp. as promising candidates for use as probiotics to promote plant vigor and activate plant defense pathways. However, the depletion of specific taxa under stress represents a mere correlation with host decline rather than a confirmed loss of essential function, necessitating experimental validation to distinguish secondary shifts from true functional causation. Furthermore, it remains unclear whether the restoration of a single keystone genus, such as *Cylindrospermum*, is sufficient to recover host immunity, or if the synergistic complexity of a full RMT is required to overcome antibiotic-driven dysbiosis. Understanding whether targeted probiotic intervention via a single cyanobacterial species can match the efficacy of a total microbiome transplant is essential for developing management techniques that improve agroecosystem resilience, reduce reliance on bactericides, and promote more sustainable, environmentally responsible agricultural practices.

This study aimed to evaluate whether targeted inoculation with filamentous nitrogen-fixing bacterium, *Cylindrospermum* sp. or RMT can mitigate antibiotic-induced rhizosphere microbiome dysbiosis in tomato plants infected with *Xanthomonas perforans*, which causes bacterial leaf spot disease. Specifically, we investigated the impact of *Cylindrospermum* inoculation or RMT on (i) rhizosphere bacterial diversity and community structure, (ii) disease severity, (iii) photosynthesis and transpiration rates, and (iv) expression of pathogenesis-and ISR-related genes.

## Materials and Methods

### Plant material, growth conditions, and experimental design

Soil was obtained from a field previously cultivated with solanaceous crops (26 m elevation, 29°38′14.064″ N, 82°21′40.6548″ W). The soil was mixed with potting soil at a 1:1 ratio to improve moisture retention. The mixed soil was subjected to a thorough study of its physiochemical characteristics, as follows: Cu^2+^(0.05 mg kg^-1^), Mn^2+^(0.57 mg kg^-1^), Ca^2+^(183.14 mg kg^-1^), Zn^2+^(0.34 mg kg^-1^), Mg^2+^(60.29 mg kg^-1^), K^+^(179.07 mg kg^-1^), P^3-^(17.39 mg kg^-1^), NH^4+^(1.60 mg kg^-1^), NO^3-^(90.91 mg kg^-1^), 2.89% of organic matter, and a pH equal to 6.71. Tomato (*Solanum lycopersicum* L. cv. ‘Ailsa Craig’) seedlings were used for all experiments. Seeds were grown in 2.7 L nursery pots in a controlled-environment greenhouse at the University of Florida plant pathology department (26 m elevation, 29°38′20.238″ N, 82°21′21.9708″ W). Temperature (25 ±2°C) and relative humidity (70-80%) were maintained within target ranges, and temporal fluctuations in these parameters were continuously monitored and recorded throughout the experiments (Figs. S1 and S2). Seedlings were maintained for three weeks prior to streptomycin treatment and pathogen inoculation and were watered every 72 h. The watering continued every 72 h after rhizosphere microbiome transfer (RMT) and *Cylindrospermum* sp. inoculum treatments.

A randomized complete block design (RCBD) with four biological replicates per treatment was used, and all plants were inoculated with a streptomycin-resistant strain of *Xanthomonas perforans*. Four treatment groups were established: (i) the control group (Ctrl), which received water only; (ii) the streptomycin group (Str), in which streptomycin was applied to induce rhizosphere dysbiosis; (iii) the streptomycin and *Cylindrospermum* sp. inoculum group (Str+CI), where plants received streptomycin followed by pathogen inoculation and *Cylindrospermum* sp., respectively; and (iv) the streptomycin and rhizosphere microbiome transplant group (Str+RMT), in which an RMT suspension was applied to plants that had received streptomycin followed by inoculation with *X. perforans*. For each treatment group, we used six replicates, and the experiment was conducted twice, yielding a total of 12 replicates per treatment.

Two hundred mL of streptomycin solution (600 mg L^−1^) were applied to the plants in the antibiotic treatment groups via soil drenching 24 h before pathogen inoculation. The control group received an equivalent volume of filtered water used for antibiotic solutions. The homogenized RMT inoculum (100 mL) and CI (100 mL) were applied via soil drench around each plant’s root zone in the Str+RMT and Str+CI group plants, respectively, 48 h post-antibiotic treatment (24 h after *X. perforans* inoculation). Thus, the control and streptomycin groups received the same volume of water (100 mL). The RMT and CI were reapplied on day 14 to the Str+RMT and Str+CI groups following the initial treatment.

### Rhizosphere soil microbe transplant (RMT) and *Cylindrospermum* sp. inoculum preparation

The rhizosphere soil microbial extraction for transplantation (RMT) adhered to the methodologies established by^6^, with certain alterations. The soil microbiome intended for transplantation into antibiotic-treated plants was obtained from the rhizospheres of three healthy (non-inoculated) tomato plants grown in mixed soil. The roots were manually agitated to eliminate excess soil, preserving only the soil intimately adhered to the roots. 100 g of root-associated soil was homogenized in three 45 s intervals, with a blending cycle of 900 mL of 0.85% NaCl solution. The resultant soil suspension was filtered through a 20-mesh (0.7 mm) screen sieve, then through a 200-mesh (0.07 mm) screen sieve, to exclude root debris and larger soil particles and obtain RMT inoculum.

A filamentous heterocyst-forming *Cylindrospermum* sp. strain LB942 was obtained from the UTEX Cultural Collection of Algae at the University of Texas at Austin and grown using BG-11^0^, a nitrogen-free medium^23^. Cultures were maintained under 12:12 light/dark (1000–2000 Lux) at 20°C. Prior to application, biomass was harvested 3 weeks after growth by centrifugation (4000 × g, 10 min) and resuspended in sterile BG-11^0^ to an OD_600_ of 0.6 to obtain a *Cylindrospermum* inoculant (CI) suspension. Filament density was confirmed microscopically using standardized hemocytometer counts (approximately 5.45 × 10^6^ cells mL^−1^ at each application).

### Pathogen inoculation and disease severity assessment

A pure *X. perforans* colony, streptomycin-resistant strain *Xp*1-6, was used to establish a liquid culture in 50 mL of lysogeny broth (LB) medium^24^, which was incubated on a shaker at 28°C and 150 rpm for 24 h to obtain pathogen cells for inoculation. The cultures were centrifuged at 10,000 × g for 10 min at room temperature (25°C) to remove the culture medium and obtain the bacterial pellet. To reach a concentration of 10^5^ cells mL^-1^, the pellet was further resuspended in phosphate-buffered saline solution to an optical density (OD_600_) of 0.1. The tomato plants were thereafter sprayed with bacterial suspension until runoff. The plants were wrapped in clear polypropylene trash bags to maintain humidity and support inoculum viability for 24 h. Starting three days post-pathogen inoculation (DPI), plants were evaluated every three days for the cumulative foliar bacterial leaf spot disease symptoms induced by the *X. perforans*, using the Standard Area Diagram (SAD) for bacterial spot on tomato leaves developed by^25^.

### Photosynthetic gas exchange measurements

Before treatments and again at 30 dpi, photosynthetic parameters were measured on the hardened, fully expanded leaves (darker green, shiny, and hard to the touch, suggesting full development and maximal photosynthetic efficiency) using a LI-6800 Portable Photosynthesis System (LI-COR Biosciences, Lincoln, NE, USA). Net CO_2_ assimilation rate (*A*, μmol CO_2_ m^−2^ s^−1^), stomatal conductance (*g*_*sw*_, mol H_2_O m^−2^ s^−1^), and transpiration rate (*E*, mmol H_2_O m^−2^ s^−1^) were recorded under controlled chamber conditions, including a photosynthetically active radiation of 1000 μmol m^−2^ s^−1^, CO_2_ concentration of 420 ppm, leaf temperature of 25°C, and relative humidity of 60%. All measurements were taken between 9:30 AM and 3:00 PM. Three measurements were collected per plant (technical replicates).

### Rhizosphere soil sampling and DNA extraction

Four weeks after the initial application of RMT and CI, rhizosphere soil was collected from each plant (six plants per treatment) to capture the full diversity and minimize variability among samples. Prior to sampling, we prepared an epiphyte removal buffer containing 6.75 g of KH_2_PO_4_, 8.75 g of K_2_HPO_4_, and 1 mL of Triton X-100 in 1 L of sterile water. The buffer was sterilized using a 0.22 μm vacuum filter. Sterile scissors were used to excise a representative segment of roots, and at least 500 mg of root tissue was transferred into a 50-mL conical tube. Thirty milliliters of epiphyte removal buffer were applied to immerse the roots, and the sample was promptly placed on dry ice, as specified by^26^. The samples were stored at −80°C thereafter. To isolate the rhizosphere soil, we followed the same method described by^27,28^.

### 16S rRNA amplicon sequencing and microbiome analysis

Amplicon library preparation and sequencing were performed by SeqCenter (Pittsburgh, PA, USA). The V3–V4 region of the 16S rRNA gene was amplified using primers 341F/806R forward: CCTACGGGDGGCWGCAG; and reverse: GACTACHVGGGTATCTAATCC^29^. Sequencing was conducted on an Illumina NextSeq2000 platform using a P1 600cyc flow cell to produce 2×300 bp paired-end reads. Raw sequencing data were processed using bcl-convert (Illumina) for quality filtering and adapter trimming. Sequences were imported into Qiime2 for further analysis. Primer sequences were removed using the cutadapt3 plugin. Sequence denoising and quality control were performed using the DADA2 plugin in QIIME2 to generate high-quality sequences. Sequencing depth was generally comparable across samples, although some variation in library size was observed. For downstream taxonomic and statistical analyses, sequences were clustered into operational taxonomic units (OTUs) using the VSEARCH5 algorithm implemented in the feature-classifier plugin of Qiime2, based on the Silva 138 reference database (99% similarity threshold)^30^. OTU tables were further processed using the SHAMAN Pasteur Institute pipeline for normalization and statistical analysis^31^ and their counts were adjusted to reflect their relative abundances within each sample. Raw read data are available at NCBI SRA under accession PRJNA1417296.

### Microbial co-occurrence network and microbiome-disease association analyses

Microbial co-occurrence networks were constructed to evaluate interaction patterns among rhizosphere microbial taxa under the control condition. Prior to network inference, low-abundance taxa were filtered to reduce noise and false correlations. Specifically, taxa with relative abundance below 0.0001 across samples were excluded. Pairwise associations among taxa were inferred using Spearman’s rank correlation coefficient. Network construction was performed using the “trans_network” function implemented in the “microeco” package. To ensure statistical robustness, only correlations with a significance level of p < 0.05 were retained. An automated threshold-selection approach based on Random Matrix Theory (COR_optimization = TRUE) was applied to determine the optimal correlation coefficient cutoff for network construction. The resulting network comprised nodes representing microbial taxa and edges representing significant pairwise correlations. Network topology metrics, including the number of nodes and edges, were subsequently used to characterize microbial interaction patterns.

To assess the relationship between microbial community composition and disease severity, the area under the disease progress curve (AUDPC) according to^32^ was calculated for each sample using disease severity measurements collected across multiple time points. The OTU table was aggregated to the genus level, and relative abundances were calculated by normalizing counts to total sequencing depth per sample. Spearman’s rank correlation analysis was performed using R (v4.2.3) to evaluate associations between the relative abundance of each genus and AUDPC values. P-values were adjusted for multiple comparisons using the Benjamini-Hochberg false discovery rate correction. Genera with significant correlations (adjusted p-value < 0.05) were considered associated with disease severity; positive correlations indicated potential disease-associated genera, and negative correlations indicated potential disease-suppressive genera.

### RNA extraction, cDNA synthesis, and RT-qPCR

Total RNA was extracted from frozen plant leaf tissue (collected 48 h post-inoculation with *X. perforans*) using TRI Reagent® Solution (Zymo Research, USA) in combination with the Direct-zol™ RNA Miniprep kit (Zymo Research, USA), following the manufacturer’s instructions with minor modifications. Briefly, approximately 50-100 mg of frozen leaf tissue was ground to a fine powder in liquid nitrogen using a pre-chilled mortar and pestle. The powdered tissue was immediately transferred to a 2 mL RNase-free microcentrifuge tube, returned to liquid nitrogen, and stored at -80°C until use. The stored samples were thawed on ice, 800 µL of TRI Reagent® was added to each, and the samples were homogenized thoroughly by vortexing for 5 m. After homogenization, we centrifuged at room temperature for 30 s at 12,000 x g. An equal volume of 99.99% ethanol was then added to the lysed supernatant collected in a new RNA-free tube, mixed thoroughly by inversion, and the mixture was transferred directly to a Direct-zol™ RNA spin column. The column was centrifuged at 12,000 × g for 30 s, and the flow-through was discarded. On-column DNase I treatment was performed using the DNase I set provided with the kit to eliminate genomic DNA contamination. The column was subsequently washed according to the manufacturer’s protocol. Total RNA was eluted in 50 µL of RNase-free water. RNA concentration and purity were assessed using a spectrophotometer (NanoDrop), and RNA integrity was verified by agarose gel electrophoresis. Extracted RNA was stored at -80°C until further use. Only RNA samples with A260/280 ratios between 1.8 and 2.1 were used for downstream cDNA synthesis and quantitative PCR (qPCR) analyses.

First-strand cDNA was synthesized using the Maxima First Strand cDNA Synthesis Kit (Thermo Scientific, USA) following the manufacturer’s guidelines. In summary, the kit components were thawed, combined, centrifuged, and kept on ice prior to use. Total RNA (500 ng) was treated with 1 µL of 10X dsDNase Buffer and 1 µL of dsDNase in a nuclease-free tube, with nuclease-free water added to a final volume of 10 µL to eliminate residual genomic DNA. The mixture was gently mixed, centrifuged, incubated at 37°C for 2 min in a water bath, and then quickly chilled on ice. Thereafter, we added 4 µL of 5X Reaction Mix, 2 µL of Maxima Enzyme Mix, and 4 µL of nuclease-free water into each reaction, resulting in a total volume of 20 µL. After gently mixing and spinning, the reactions were incubated at 25°C (room temperature) for 10 min, then at 50°C for 15 min in a water bath. Enzyme activity was inactivated by heating at 85°C for 5 min in a water bath. The resultant cDNA for quantitative PCR (qPCR) was immediately preserved at -80°C until use.

Quantitative real-time PCR (qPCR) was conducted utilizing a CFX Opus 96 Real-Time PCR System (Bio-Rad Laboratories, Hercules, CA, USA). Reactions were performed in a total volume of 20 µL, containing 2 µL of cDNA template, 10 µL of 2X SYBR Green master mix, 0.2 µL of each gene-specific primer, and nuclease-free water to bring the final volume to 20 µL. All reactions were conducted in technical triplicate. Thermal cycling parameters included an initial denaturation at 95°C for 15 m, followed by 40 cycles including denaturation at 95°C for 15 s, annealing at 59°C for 30 s, and extension at 72°C for 30 s. Fluorescence data were acquired following the conclusion of each annealing phase. A melt curve study was conducted post-amplification by incrementally raising the temperature from 65°C to 95°C in 0.5°C intervals, while continuously acquiring fluorescence to verify amplification specificity. Information concerning the qPCR primers is provided in Table 1. Gene expression levels were determined via the 2^−ΔΔCt^ method^33^, with normalization to the chosen reference genes. Primer specificity and amplification efficacy were confirmed before analysis.

**Table 1.**
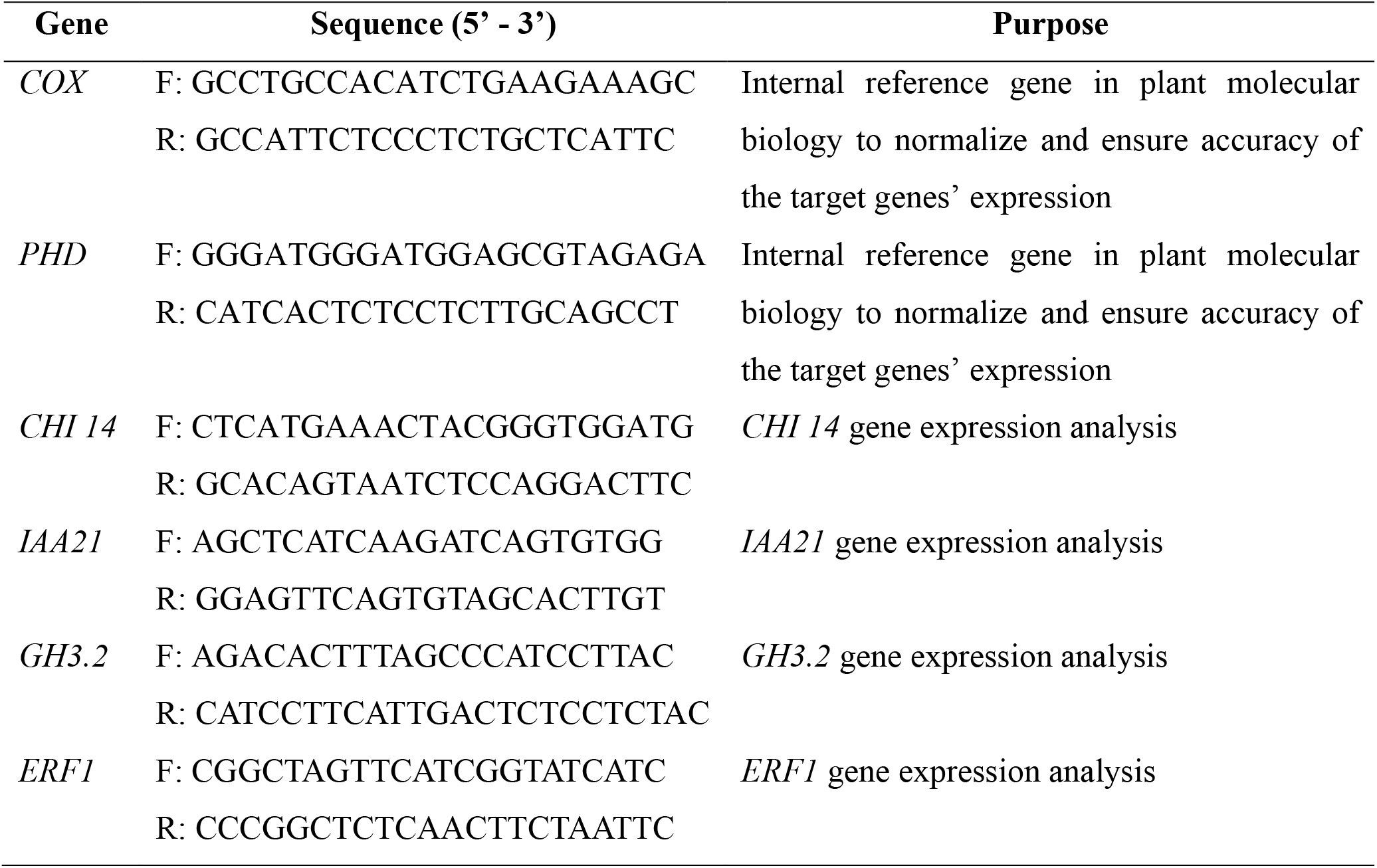
Primers used in qPCR analysis.

### Statistical analysis

Data analyses were performed in R Studio (v4.3.1). Disease severity and photosynthetic parameters from the two experiments were analyzed using Kruskal-Wallis one-way ANOVA followed by Tukey’s HSD test (p < 0.05). Graphs were generated using *ggplot2* package. SHAMAN, an online bioinformatics tool for processing raw reads in targeted metagenomics (https://shaman.pasteur.fr/), was used to assess bacterial 16S rRNA gene sequences using statistical methods. Bar plots of the microbial community and Shannon curves were generated to characterize rhizosphere diversity. Principal coordinates analysis (PCoA) was performed on normalized relative abundance data to elucidate trends in microbial community composition. The Bray-Curtis dissimilarity metric was used to measure variation in community composition between samples.

Shannon Index values were computed to assess alpha diversity across treatments. Statistical differences in community composition were evaluated using permutational multivariate analysis of variance (PERMANOVA) based on Bray-Curtis distances, and Wilcoxon tests were used for pairwise comparisons of diversity indices (*P* < 0.05). The Shannon and Inverse Simpson diversity indices were computed using the default values established by^31^. Differences in Cyanobacteria and *Cylindrospermum* abundance among treatments were evaluated using the Kruskal-Wallis test, followed by Dunn’s post-hoc test with Benjamini-Hochberg correction for multiple comparisons. Statistical analyses were conducted in R (v4.2.3). All experiments in this study were repeated to confirm the results.

## Results

### *Cylindrospermum* sp. inoculum and RMT reduced disease progression under antibiotic-induced dysbiosis

To induce dysbiosis, streptomycin and water (control) were initially applied to the rhizosphere of tomato plants. 24 h later, a streptomycin-resistant strain of the foliar pathogen *X. perforans* was sprayed onto the upper portion of the plants. The RMT slurry and *Cylindrospermum* inoculum (CI) were applied to the rhizosphere at the root zone of each Str+RMT and Str+CI set of plants, respectively, 48 h after antibiotic treatment (thereby, 24 h after *X. perforans* spray). The streptomycin application resulted in increased disease severity at 12 and 15 days post-probiotic application, with increases of 18% and 17.8%, respectively, compared with control plants (Fig. 1A). Plants treated with RMT or CI after Str application showed a significant reduction in disease severity from Day 6 onward compared to untreated plants. These differences were visually corroborated by pronounced foliar symptoms at 30 days after the first RMT and CI applications (Fig. 1B).

**Fig. 1.**
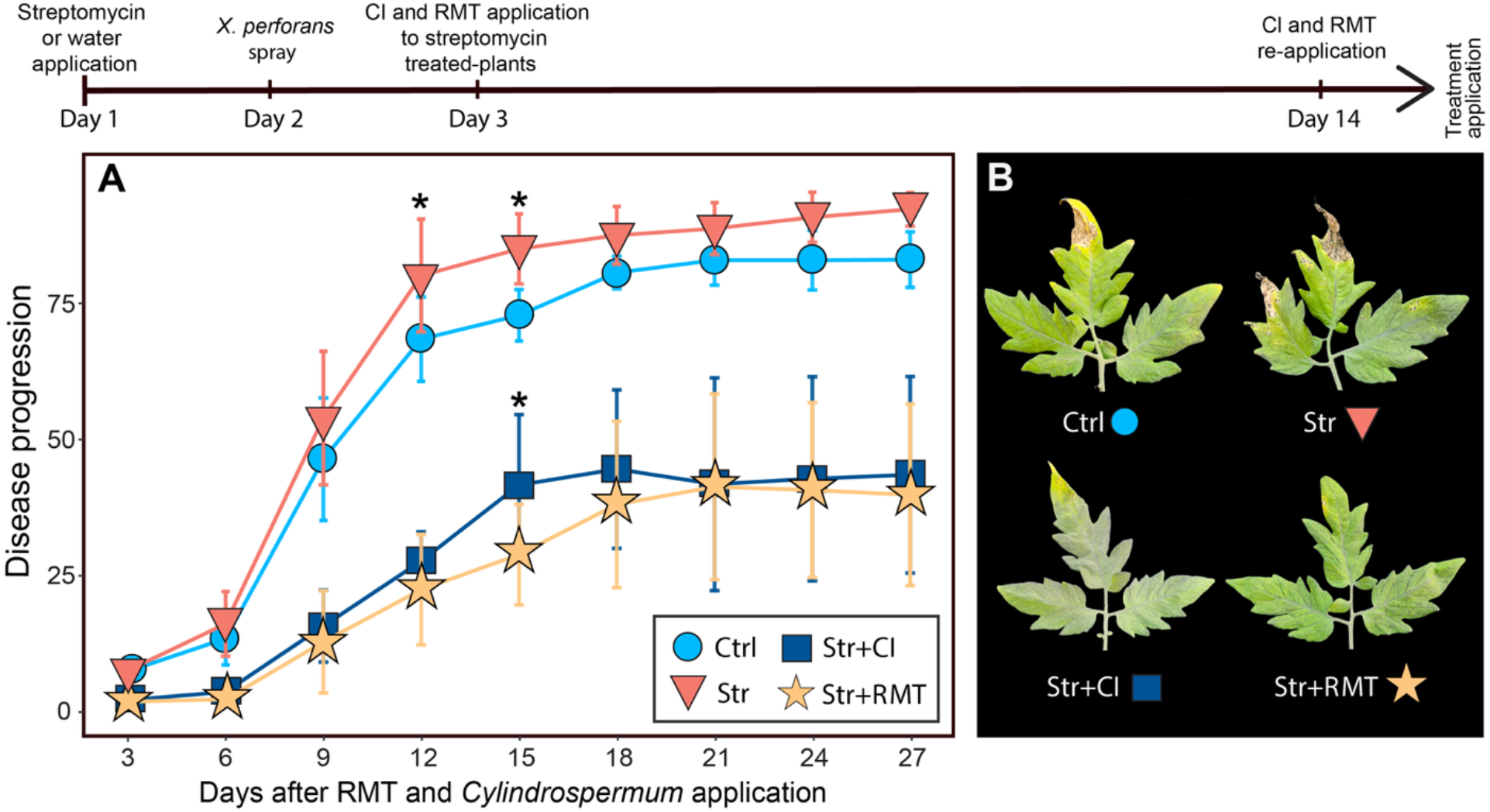
Effects of streptomycin-induced dysbiosis and RMT and *Cylindrospermum* sp. inocula on disease severity and foliar symptom development in tomato. Plant rhizosphere was treated with 0.6 g L^-1^ streptomycin or water (control), followed by spray inoculation with *X. perforans* (OD_600_ = 0.1, approximately 10^5^ cells mL^-1^) after 24 h. Two groups of plants that received streptomycin and *X. perforans* were administered rhizosphere soil microbiome transplants (RMT) from uninfected (healthy) plant donors or *Cylindrospermum* sp. inoculum (CI) 48 h post-streptomycin administration. (A) Disease severity progression following RMT and CI applications, showing increased severity at 12 and 15 days post-RMT/CI applications compared with control plants. Data represent mean values ±SE. Average of two experiments with six replicates for each treatment (n=12). The asterisk (*) indicates statistically significant differences between Str-treated plants and the Ctrl plants (P<0.01). Although additional significant differences were detected between non-inoculated treatments (Ctrl and Str) and the microbial inoculation treatments (Str+CI and Str+RMT) at several time points, these comparisons are not individually marked in the figure to maintain clarity. Complete statistical results are provided in Supplementary Table S1. (B) Representative images of leaflet symptoms at 30 days post-streptomycin application (leaflets were collected on the 8^th^ node of each plant per treatment), illustrating enhanced disease symptom development under antibiotic-induced dysbiosis and differential responses to RMT and CI treatments.

### Rhizosphere bacterial assemblages in response to RMT and CI treatments

To identify key bacterial groups that shape rhizosphere community structure and their potential responses to treatment conditions, we examined the most abundant taxa. Figs. 2A and 2B illustrate the twelve and ten most prevalent phyla and genera, respectively, in the rhizospheres of tomato plants after 30 days post-pathogen inoculation. Given our focus on the effects of CI and RMT following streptomycin application, we conducted pairwise comparisons of median abundances within the cyanobacteria phylum to assess treatment-specific responses. Compared with the control, streptomycin alone reduced cyanobacteria relative abundance by 9.4%, whereas Str+CI increased the abundance by 42.5%. The Str+RMT treatment caused only a minor decrease (2.5%) compared with the control. When treatments were compared directly, Str+CI showed substantially higher abundance than both streptomycin alone (57.3% increase) and Str+RMT (46.3% increase). In contrast, differences between streptomycin and Str+RMT were small (7.6%), indicating limited divergence between these treatments.

**Fig. 2.**
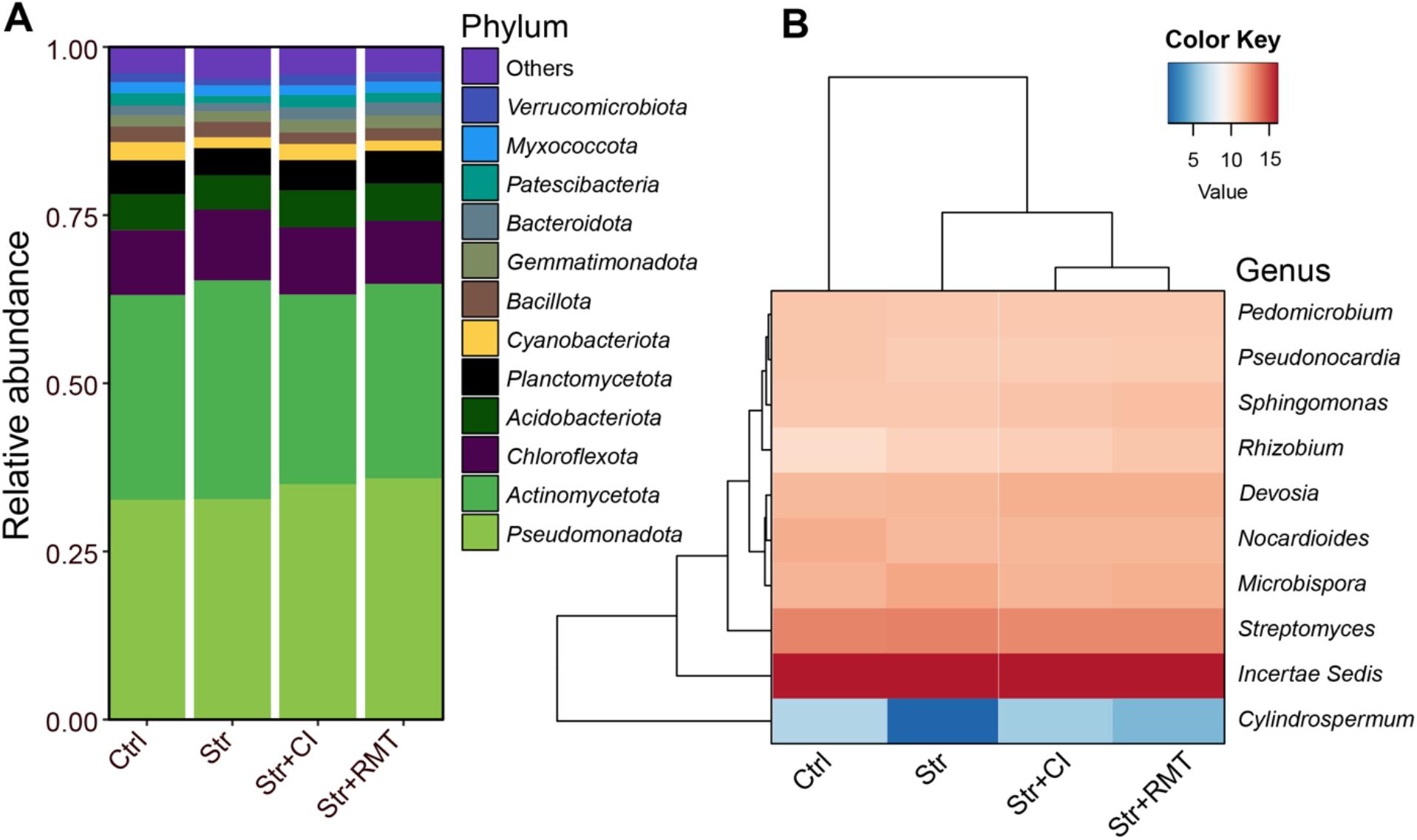
Relative abundance of rhizosphere microbial communities at the phylum (A) and genus (B) levels across all treatment groups. Bar plots (A) and a heatmap (B) show the mean proportional abundance of dominant taxa (two experiments, six replicates per treatment, n=12), illustrating shifts in community structure 30 days following *X. perforans* infection, as well as the extent of microbiome recovery in response to *Cylindrospermum* sp. inoculation and rhizosphere microbiome transplant treatments. Low-abundance taxa are grouped as “Other.” Color Key represents row-scaled Z-scores of log-transformed relative abundances, with red indicating higher and blue indicating lower abundance relative to the mean across samples.

Genus-level analysis revealed treatment-specific responses of *Cylindrospermum* following streptomycin application. Pairwise comparisons of median *Cylindrospermum* abundance indicated that streptomycin significantly reduced *Cylindrospermum* relative abundance when compared with the control, whereas the addition of CI (Str+CI) resulted in a marked increase in its abundance. In contrast, the Str+RMT treatment caused only modest changes in *Cylindrospermum* abundance compared with streptomycin, indicating limited impact on *Cylindrospermum* recovery. Direct comparisons among treatments further showed that *Cylindrospermum* relative abundance was substantially higher under Str+CI than under both streptomycin and Str+RMT, suggesting that targeted inoculation more strongly influenced *Cylindrospermum* responses than microbiome transplantation.

Alpha diversity metrics did not differ significantly among treatments (Fig. 3). Shannon richness diversity showed a marginal trend toward treatment effects (Kruskal-Wallis, p = 0.052), with slightly higher median values in the Str+CI and Str+RMT treatments than in the control and streptomycin alone (Fig. 3A). Similarly, microbial evenness did not vary significantly among treatments (Kruskal-Wallis, p = 0.21; Fig. 3B), indicating comparable community evenness across all groups. However, the streptomycin treatment alone reduced the evenness compared to the control. Principal coordinates analysis (PCoA) based on Bray-Curtis dissimilarity revealed overlap among treatment groups, with no clear separation along the first two axes, which together explained 69.0% of the total variance (Axis 1: 52.97%; Axis 2: 16.03%) (Fig. 3C).

**Fig. 3.**
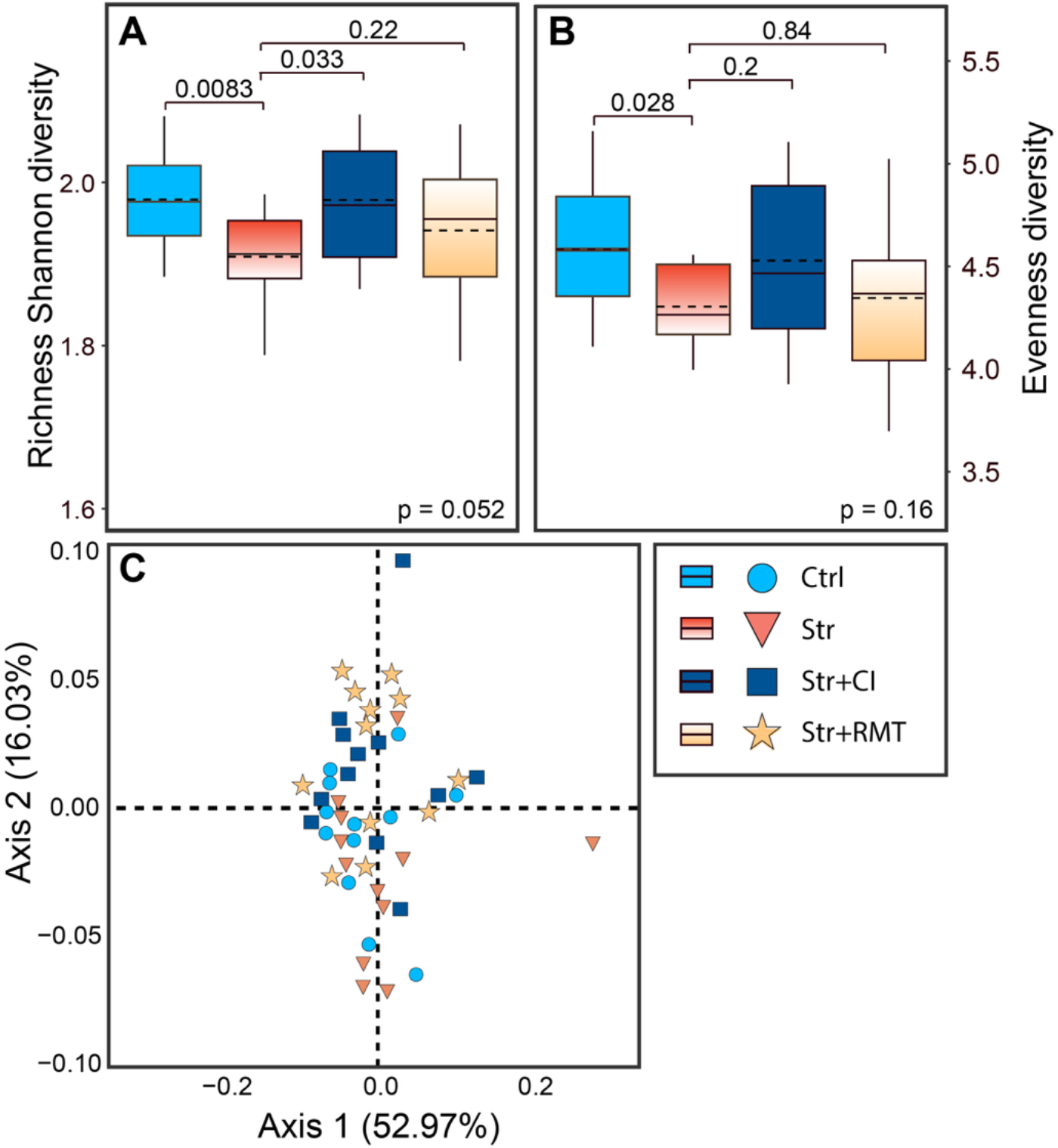
Alpha and beta diversity of bacterial communities across treatments. (A) Shannon diversity and (B) Evenness diversity across control (Ctrl), streptomycin (Str), streptomycin plus *Cylindrospermum* sp. (Str+CI), and streptomycin plus rhizosphere microbiome transplant (Str+RMT) treatments. Boxes represent the median and interquartile range, with points indicating individual biological replicates. Differences among treatments were assessed using the Kruskal-Wallis test. (C) Principal coordinates analysis (PCoA) based on Bray-Curtis dissimilarity showing bacterial community composition across treatments. Percentages on axes indicate the variance explained by each axis. Means of two experiments, six replicates per treatment (n=12).

### Treatment-dependent microbial network architecture

Co-occurrence network analysis indicates that Str treatment results in significant reshaping of the rhizosphere microbial community network architecture (Fig. 4). The control network, characterized by high connectivity (1,104 nodes, 4,090 edges) and mostly positive interactions (94.77%), indicates a stable community (Fig. 4A). In contrast, Str treatment reduced connectivity (1,031 nodes, 2,870 edges) and increased negative interactions (10.77%) (Fig. 4B), suggesting microbial destabilization. The Str+CI treatment showed intermediate complexity (1,081 nodes, 2,955 edges) with a further increase in negative correlations (15.47%), indicative of potential competitive interactions (Fig. 4C). Since the treatment involved the introduction of *Cylindrospermum* sp. following antibiotic perturbation, the elevated negative associations likely reflect competitive restructuring between the introduced inoculum and resident taxa during microbiome reassembly. Str+RMT resulted in a network comparable to the control regarding nodes and edges numbers (1,101 nodes, 4,220 edges) but demonstrated the highest rate of negative interactions (25.64%) (Fig 4D). This pattern indicates extensive restructuring of microbial interactions, likely reflecting competitive dynamics between introduced and native taxa during microbiome reassembly rather than restoration of the native bacterial community state.

**Fig. 4.**
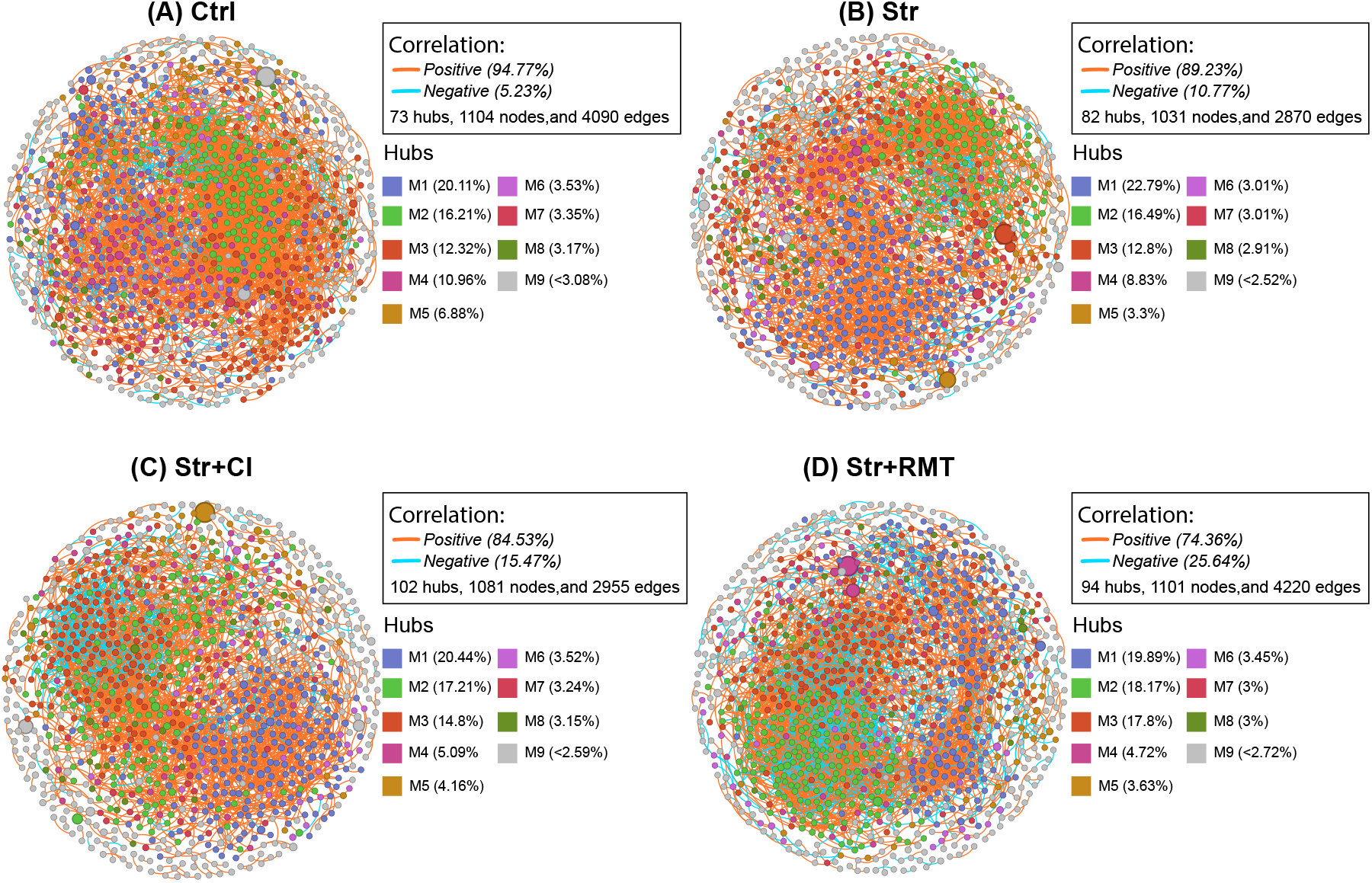
Co-occurrence networks of rhizosphere bacterial communities across treatments. Networks were constructed based on Spearman correlations among OTUs for (A) Control, (B) Streptomycin, (C) Streptomycin + CI, and (D) Streptomycin + RMT treatments. Nodes represent bacterial taxa and edges represent significant correlations (|ρ| ≥ threshold, adjusted p < 0.05). Edge colors indicate positive (red) and negative (blue) associations, and node size reflects connectivity. Hub taxa (M1-M9) represent highly connected modules within each network, with their relative contributions indicated as percentages.

### Plant phenotype (disease)-microbiome associations align with treatment-driven network structure

Integration of disease severity and microbiome composition revealed strong links between specific bacterial taxa and disease outcomes across treatments. Disease severity, quantified as AUDPC, varied markedly among treatments, with streptomycin-treated plants exhibiting higher disease levels, while microbiome restoration treatments (Str+CI and Str+RMT) reduced disease severity relative to antibiotic-only conditions (Fig. 5A). Correlation analysis further identified key genera associated with disease progression: several were positively correlated with AUDPC, whereas others showed negative associations, suggesting potential disease-promoting and disease-suppressive roles, respectively (Fig. 5C). Notably, the relative abundance of *Cylindrospermum* was negatively correlated with disease severity (ρ = -0.38, p = 0.0083; Fig. 5B), indicating that increased representation of this taxon is associated with reduced disease and depletion of this taxon with higher disease severity. Antibiotic disturbance altered both community composition and interaction structure, while subsequent microbial interventions promoted distinct modes of community reassembly, ranging from competitive interactions following targeted inoculation to large-scale restructuring following microbiome transplantation.

**Fig. 5.**
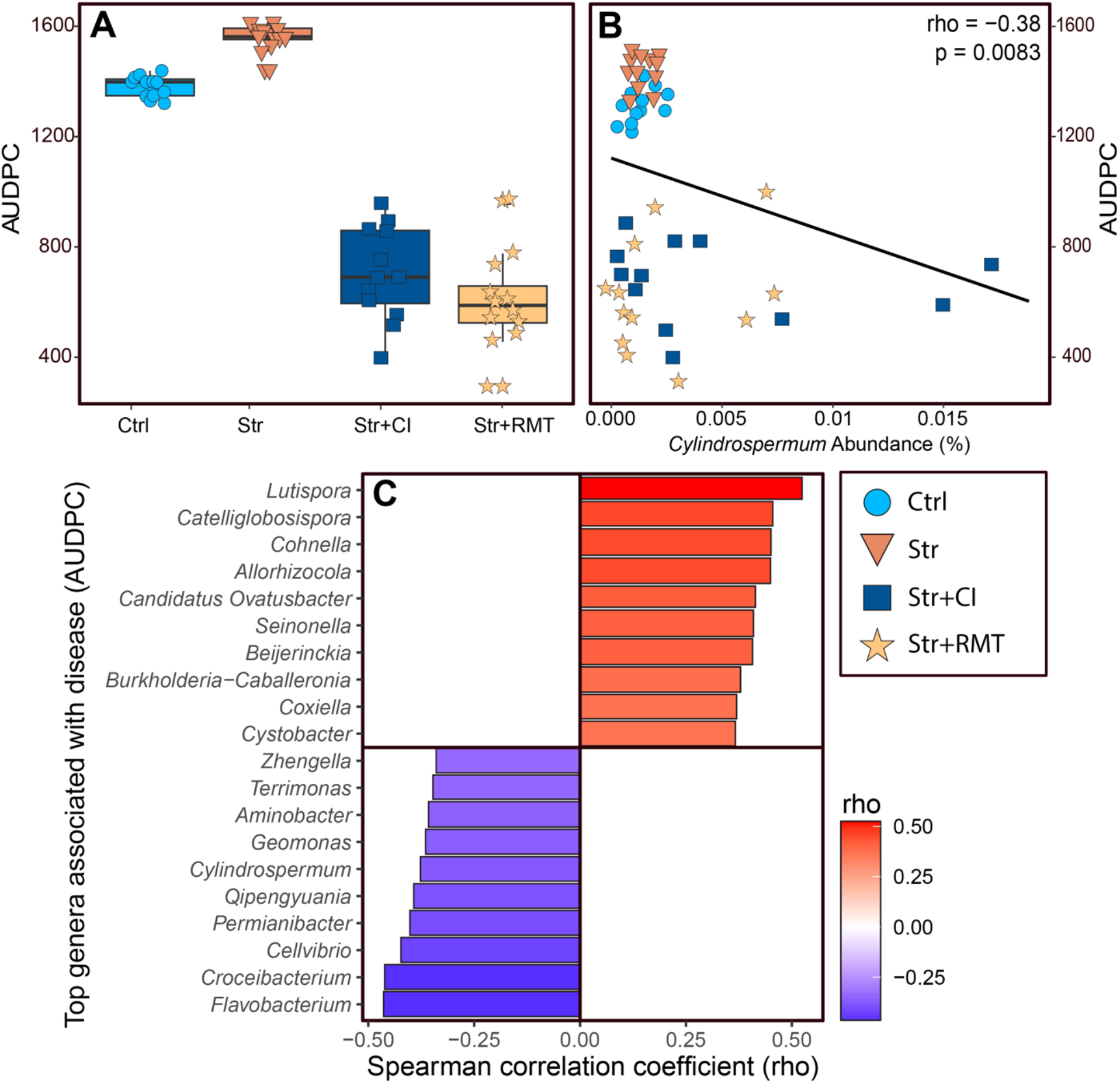
Integration of disease severity and microbiome composition across treatments. (A) Disease severity, measured as the area under the disease progress curve (AUDPC), across treatments (Control, Streptomycin, Streptomycin + *Cylindrospermum* sp. (CI), and Streptomycin + RMT). Points represent individual samples, and boxplots summarize the distribution within each treatment. (B) Relationship between the relative abundance of *Cylindrospermum* sp. and disease severity (AUDPC). Each point represents a sample colored by treatment. A significant negative correlation was observed (Spearman’s ρ = -0.38, p = 0.0083), indicating that higher *Cylindrospermum* abundance is associated with reduced disease severity. (C) Top bacterial genera associated with disease severity based on Spearman correlation analysis. Positive correlations (red) indicate taxa associated with increased disease severity, whereas negative correlations (blue) indicate taxa associated with reduced disease severity. Only the most strongly associated genera are shown, ranked by their correlation coefficients.

### Photosynthetic and transpiration responses to *Cylindrospermum* sp. inoculation

Two time points were evaluated to examine physiological characteristics associated with streptomycin treatment of the rhizosphere. The initial measurements were taken a day before antibiotic application, when the plants were healthy and thus not infected with *X. perforans*. The second measure occurred on day 30 following antibiotic treatment. Our results showed that, under the conditions of the experiment, despite pronounced differences in disease severity among treatments, net assimilation and transpiration rates remained statistically unchanged (P > 0.05), suggesting that disease progression under antibiotic-induced dysbiosis occurred independently of measurable alterations in leaf gas exchange (Fig. 6).

**Fig. 6.**
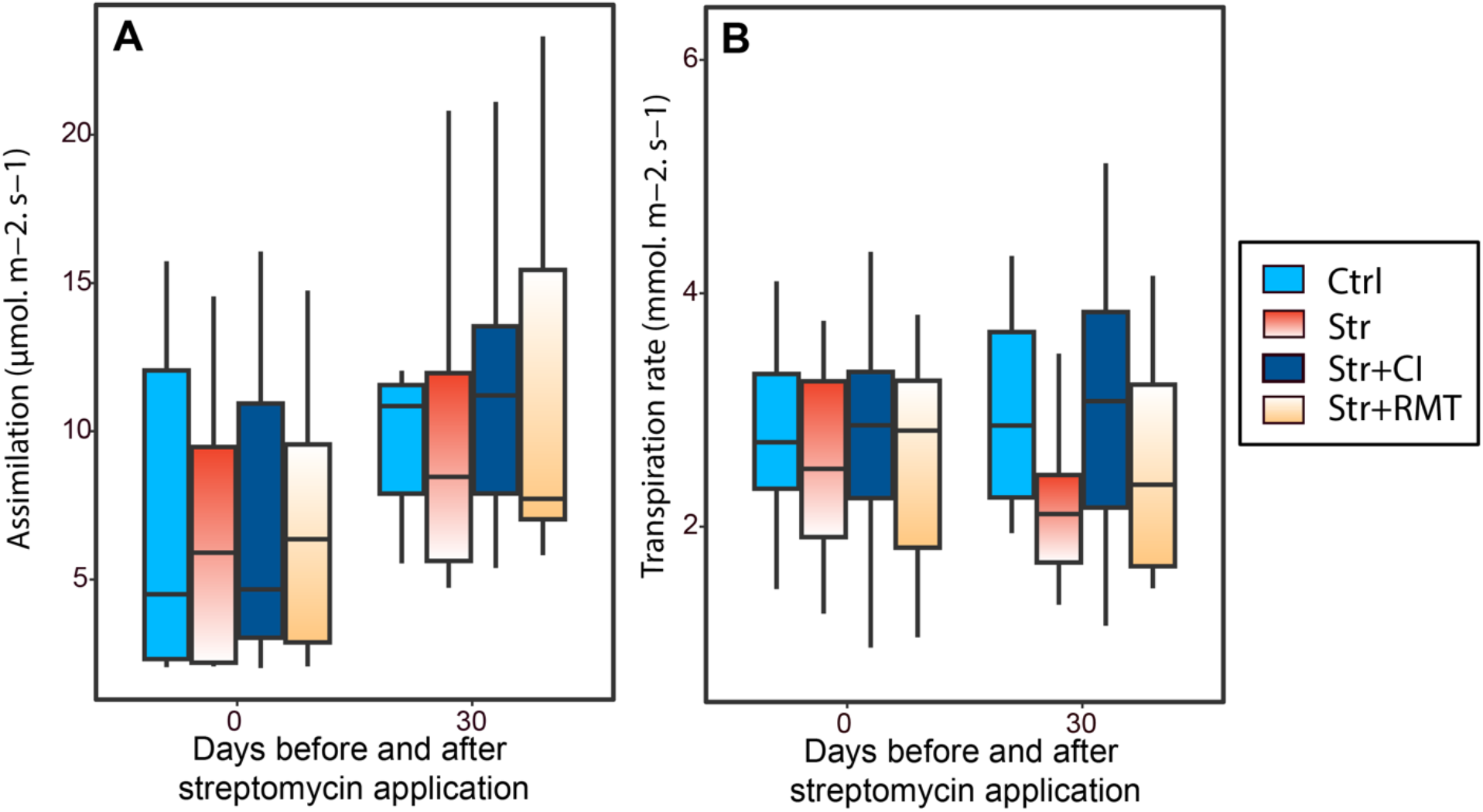
Photosynthetic and transpiration responses to antibiotic treatment and microbial inoculation. (A) net photosynthetic rate and (B) transpiration rate measured before treatment (Day 0) and 30 days after antibiotic application (corresponding to 28 days after RMT and *Cylindrospermum* sp. inoculation) across the four treatments. Boxes represent the interquartile range (IQR), the center line indicates the median, and whiskers indicate data dispersion (data of two experiments and six replicates per treatment). No significant differences were detected among treatments at either time point (P > 0.05). Each box represents the mean of 12 replicates.

### *Cylindrospermum* sp. inoculum- and RMT-mediated regulation of plant defense genes

To corroborate the transcriptomic findings from our previous study^27^, the expression levels of two defense-related genes (*Chi14* and *ERF1*) and two auxin-responsive genes (*Gh3*.*2* and *IAA21*) were measured by qPCR. In line with previous RNA-seq findings^27^, *ERF1* expression was upregulated following streptomycin treatment compared with the control group. *IAA21* was downregulated in the presence of streptomycin, consistent with transcriptome trends. Nonetheless, qPCR results for *Chi14* and *Gh3*.*2* did not fully align with RNA-seq trends, as *Chi14* expression decreased while *Gh3*.*2* expression increased following streptomycin administration (Fig. 7). These discrepancies indicate gene-specific heterogeneity between transcriptomic and targeted expression measurements. Microbiome manipulation interventions (Str+CI and Str+RMT) mitigated the streptomycin-induced elevation of *ERF1* and reduced *Gh3*.*2* expression to control levels. Conversely, *Chi14* expression remained relatively low throughout treatments.

**Fig. 7.**
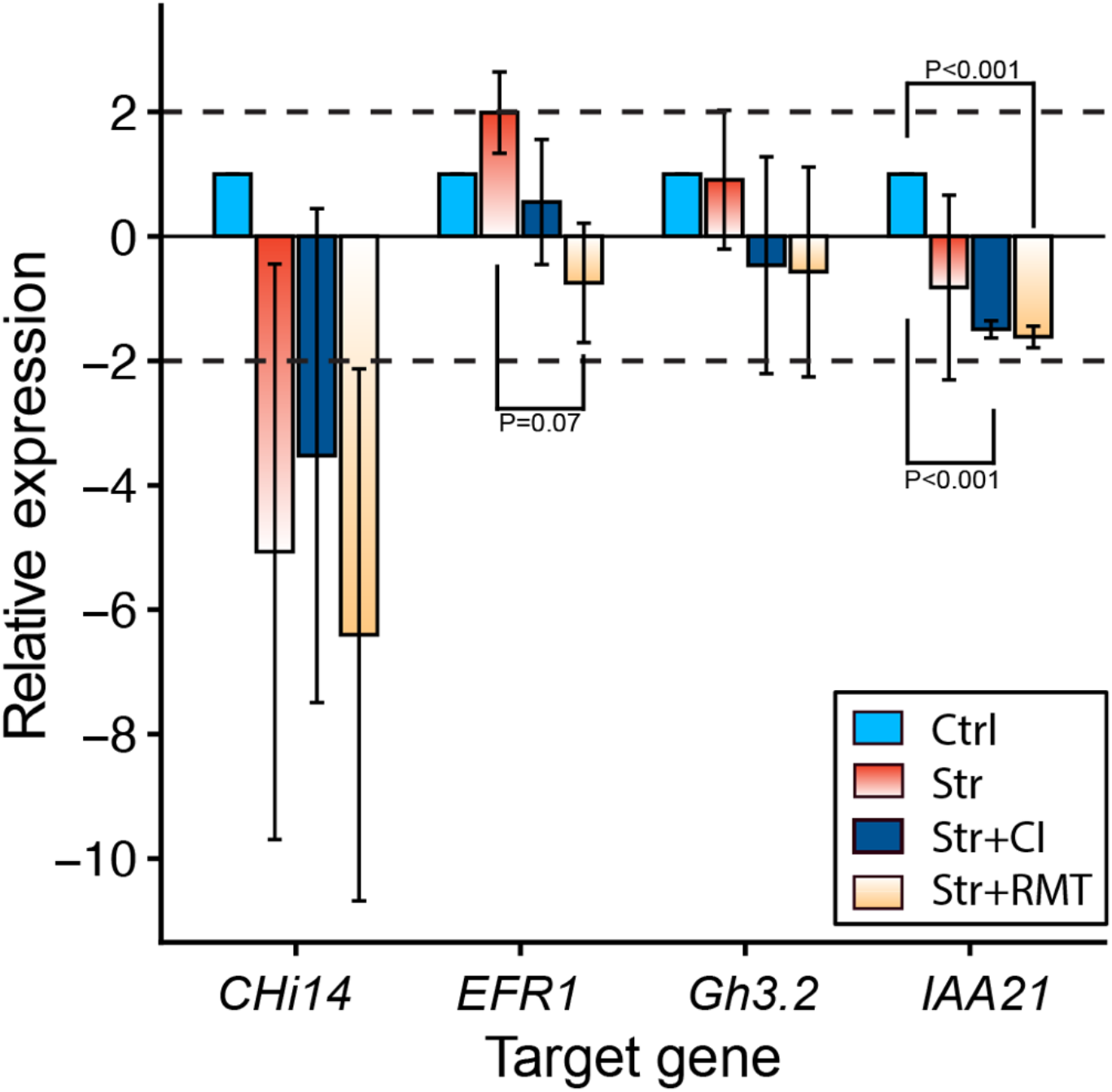
Effects of *Cylindrospermum* sp. (CI) and rhizosphere microbiome transplantation (RMT) on gene expression related to defense and auxin in tomato leaves after streptomycin treatment. qPCR analysis indicated that streptomycin increased the expression of defense-related genes *Chi14* and *ERF1*, while decreasing the expression of auxin-responsive genes *Gh3*.*2* and *IAA21* compared to the control. Both CI and RMT treatments reduced the induction of defense genes and partially normalized auxin-responsive gene expression back to control levels. Values represent mean ±SE (n = 3 biological replicates and 3 technical replicates each). Dashed lines indicate ±2 regulation threshold.

## Discussion

The rhizosphere microbiome is a critical determinant of plant fitness, governing essential processes including nutrient cycling, immune priming, and pathogen suppression. Consequently, the disruption of microbial equilibrium, often termed dysbiosis, can impair these beneficial functions and negatively affect plant physiology, reducing the plant’s ability to cope with stress, even in the absence of direct pathogen pressure^34,35^. In this study, we demonstrated that chemically induced dysbiosis significantly increased plant vulnerability to *Xanthomonas perforans* infection, an effect characterized by depletion of the indigenous cyanobacterium *Cylindrospermum*. These findings reinforce the emerging concept that antibiotic exposure can indirectly exacerbate disease severity by destabilizing the native rhizosphere architecture^28^. Notably, we found that this dysbiotic state is reversible. The reintroduction of a *Cylindrospermum* sp. (CI) alone was sufficient to delay disease progression, performing comparably to, and in some respects more efficiently than, the wholesale restoration of the community via Rhizosphere Microbiome Transplant (RMT). While both interventions facilitated recovery of the CI population, the targeted approach with CI achieved more robust colonization, suggesting that specific keystone taxon may be more pivotal to functional restoration than overall community richness. These results point toward a probiotic approach to microbiome management: from the complex ‘rebuilding’ of communities to the strategic ‘repair’ of specific functional gaps.

Co-occurrence network analysis revealed that streptomycin application reduced overall connectivity and increased negative associations, a shift consistent with the disruption of cooperative microbial relationships and the destabilization of community structure. Such structural transitions are characteristic of dysbiosis, where fragmented interaction networks often impair key ecosystem functions linked to plant fitness^36,37^. Our findings demonstrated that microbiome interventions triggered distinct modes of community reassembly rather than a simple reversion to the baseline state. Specifically, targeted inoculation with *Cylindrospermum* (Str+CI) increased negative interactions while maintaining intermediate network complexity, suggesting competitive dynamics between the introduced inoculum and resident taxa during reassembly. In contrast, rhizosphere microbiome transplantation (Str+RMT) produced a highly connected but interaction-rich network with the highest proportion of negative associations, indicative of extensive restructuring driven by competition among coexisting microbial populations. These contrasting patterns highlight that microbiome restoration strategies can lead to alternative network configurations rather than a return to the original state.

Our genus-level analyses highlight a contrast in restoration efficiency: while targeted inoculation (Str+CI) restored *Cylindrospermum* relative abundance to near-control levels, RMT provided only a modest recovery despite introducing of a high-diversity microbial assemblage. This discrepancy suggests that the successful suppression of *X. perforans* in this system may not require whole community replacement, but rather the strategic reinstatement of specific functional keystones. While previous studies have emphasized the success of transferring entire suppressive microbiome^6,38,39^, our results align more closely with the emerging framework of precision microbiome engineering efforts, such as synthetic microbial communities (SynComs)^40-42^.

Integration of microbiome composition with disease severity further identified specific taxa associated with disease outcomes, with several genera exhibiting positive or negative correlations with AUDPC. Similar patterns have been reported in plant disease systems, where specific microbial taxa are enriched or depleted in association with plant phenotype and can predict disease progression status^43,44^. Notably, the relative abundance of the *Cylindrospermum* genus was inversely correlated to disease severity, suggesting a potential role in host resilience. Previous studies have shown that plant-associated microbiomes contribute directly to disease suppression and host fitness, with specific taxa playing key functional roles within these communities^45,46^. These associations occur within microbial communities undergoing treatment-driven reorganization, indicating that disease outcomes are shaped by both microbial composition and interaction structure rather than by individual taxa alone.

Comparison of the Str+CI and Str+RMT rhizosphere communities showed that *Cylindrospermum* recovery was a specific response to targeted inoculation rather than an incidental restoration. The fact that Str+CI-treated plants exhibited disease suppression comparable to Str+RMT-treated plants suggest that this genus or its specific functions contribute to host resilience. This finding aligns with emerging evidence that specific microbial taxa can exert disproportionate influence on plant health outcomes, even when global community structure appears largely unchanged^47,48^. This outcome underscores a key limitation of RMT: the failure of introduced communities to recover specific functional keystones when faced with the ecological barriers of a disturbed rhizosphere^49^. Such findings align with ecological theory, which suggests that the targeted restoration of key functional nodes within a network can have outsized effects on system behavior, even if overall diversity metrics remain stable^50,51^.

Despite treatment-specific differences in disease severity and *Cylindrospermum* abundance, alpha- and beta-diversity analyses revealed no strong restructuring of the overall bacterial community across treatments. While streptomycin-induced dysbiosis initially reduced richness and evenness, and targeted CI application restored richness to control levels, overall diversity indices remained largely comparable across treatments, with PCoA analyses showing substantial overlap among groups. This apparent disconnect highlights a critical limitation in relying solely on broad diversity metrics to assess microbiome-plant interactions. Complex yet functionally meaningful changes in specific keystone taxa or metabolic pathways can drive plant health outcomes without producing detectable differences in overall community structure^52-54^. In this context, genus-level responses of *Cylindrospermum* provide more mechanistic insight than community-wide diversity indices, emphasizing the need for targeted taxonomic and functional analyses when evaluating microbiome-plant interactions.

Interestingly, differences in disease severity were not accompanied by measurable changes in photosynthetic rate or transpiration. Net assimilation and transpiration remained statistically unchanged across treatments, even at 30 days post-antibiotic application. This apparent decoupling between symptom development and leaf gas exchange has been reported previously, as foliar disease impacts on photosynthesis and transpiration are not universally proportional to visual severity and can vary substantially across pathosystems and contexts ^55^. This suggests that disease progression under antibiotic-induced dysbiosis occurred independently of detectable alterations in leaf gas exchange. However, because leaf gas exchange was measured only at the end of the experiment, after disease severity had largely stabilized, it is possible that transient physiological differences occurred earlier during disease development and were not captured in our measurements. This physiological stability arose despite significant transcriptional reprogramming in defense and hormone-related pathways. Streptomycin treatment significantly increased the ethylene-responsive transcription factor *ERF1* and the auxin-responsive gene *Gh3*.*2*, while downregulating defense marker *Chi14* and the auxin-responsive gene *IAA21*, indicating activation of immune signaling and a reconfiguration of hormonal balance. *ERF1* serves as a key integrator of ethylene and jasmonate signaling pathways, modulating downstream defense genes in response to bacterial challenges^56-58^. The simultaneous suppression of auxin-responsive genes aligns with the established antagonistic interaction between auxin and defense signals during immunological activation^59^.

## Conclusions and Future Directions

This study demonstrates that antibiotic-induced rhizosphere dysbiosis can exacerbate bacterial disease severity in tomato, a process characterized by the depletion of the keystone cyanobacterium *Cylindrospermum* spp. Species of We showed that this dysbiotic state is reversible; targeted inoculation with one strain of *Cylindrospermum* alone was sufficient to delay disease progression, performing with higher colonization efficiency than wholesale community restoration via RMT. These results support a shift in microbiome management toward the strategic “repair” of functional gaps rather than complex community “rebuilding”. Our findings provide mechanistic insight by connecting disease outcomes to host transcriptional reprogramming. Streptomycin-induced dysbiosis triggered a significant reconfiguration of immune and hormone signaling, through the upregulation of *ERF1* and modulation of *Chi14, Gh3*.*2*, and *IAA21*, indicating a reconfiguration of ethylene-mediated defense and auxin signaling pathways under dysbiosis. *Cylindrospermum* inoculation and rhizosphere microbiome transfer significantly reduced the overactivation of defense genes and partially restored auxin-related gene expression patterns, indicating that microbial community composition influences host immune-hormonal networks during pathogen challenges. Ultimately, this work underscores the need to evaluate the indirect ecological costs of antibiotic use and provides a framework for precision microbiome engineering in in sustainable agricultural systems.

## Acknowledgments

This work was supported by the USDA National Institute of Food and Agriculture (NIFA) project no. 2022-68015-36721 and by the Research Capacity Fund (Hatch) program, project award no. 7010682.

